# I Am A Scientist: Overcoming implicit bias in STEM through explicit representations of diversity in lectures

**DOI:** 10.1101/2022.06.23.497359

**Authors:** Dominic Henri, Kirra Coates, Katharine Hubbard

## Abstract

The lack of diversity in Science, Technology, Engineering and Mathematics (STEM) is a significant issue for the sector. Many organisations and educators have identified lack of representation of historically marginalised groups within teaching materials as a potential barrier to students feeling that a STEM career is something that they can aspire to. A key barrier to addressing the issue is providing accessible and effective evidence-based approaches for educators to implement. In this study, we explore the potential for adapting presentation slides within lectures to ‘humanise’ the scientists involved, presenting their full names and photographs alongside a Harvard style reference. We adopt a questionnaire based methodology, and survey 161 undergraduates and postgraduates at a UK civic university. We first establish that students project implicit biases onto a hypothetical reference, with over 50% of students assuming the scientist was male and Western. We then explore what students think of the ‘humanised’ slide design, concluding that most students see it as good pedagogical practice, with some students positively changing their perceptions about diversity in science. We were unable to compare responses by ethnicity, but find evidence that female and non-binary students are more likely to see this as good pedagogical practice, perhaps reflecting white male fragility in being exposed to initiatives designed to highlight diversity. We conclude that humanised powerpoint slides are a potentially effective tool to highlight diversity of scientists within existing research-led teaching, but highlight that this is only a small intervention that needs to sit alongside more substantive work to address the lack of diversity in STEM.

## Introduction

It has long been recognised that there is a lack of diversity within Science, Technology, Engineering and Mathematics (STEM) compared to the general population, and that this lack of diversity represents a loss of talent within the sector. A recent report into the UK STEM workforce identifies that 65% of STEM professionals are white men, and that women are particularly underrepresented [1]. Other measures of success in science show similar bias; 90% of Fellows of the UK Royal Society are male [2]. Researchers from Black, Asian and Ethnic Minority backgrounds are less likely to be awarded governmental research funding, and are funded with smaller grants [3]. Lesbian, Gay, Bisexual, Trans and Queer (LGBTQ+) students are less likely to complete a STEM qualification than their heterosexual peers [4]. Diverse research teams have been found to have higher productivity, and are more likely to focus on under-researched topics [5–7]. Several studies have shown that men are more likely to be hired for technical positions even when objective evidence is provided that the female candidate is equally capable [8–10]. Loss of talented individuals from the STEM workforce is a matter of significant concern, and proactive measures are increasingly being put in place to attract and retain diverse members of the scientific community.

### A white male Western bias persists throughout formal STEM education

In order to become a STEM professional, an individual must view a technical career as something that they want and can realistically achieve. They must also be able to “see themselves” working in STEM, and adopting a scientific identity [11,12]. Assumptions about ‘who’ can be a scientist are established at a young age, and persist throughout education. The ‘Draw A Scientist Test’ (DAST) is a commonly used methodology to explore children’s conceptions of scientists, used internationally for at least 50 years [13–15]. Children typically draw scientists as male, wearing a lab coat, and performing chemistry related tasks, although more recent studies have had a less masculine bias [13,15]. Older children are more likely to draw male scientists [15], suggesting that masculine stereotypes of scientists are reinforced through formal school education. Teaching resources may reinforce this perception that scientists are white and male. For example, a recent analysis of high-school level chemistry textbooks used in three different countries found a significant male bias in the scientists presented [16]. Representations of men tended to be as active scientists, whereas women were more likely to be presented in non-scientific contexts such as domestic settings [16]. Some evidence suggests that there has been little progress in improving representation within textbooks since earlier analyses in the 1970s and 1990s [17,18]. Other evidence suggests that female representation within textbooks is increasing in line with the proportion of females within the field [19]. However, studies generally agree that the ethnicity, gender, disability status, nationality, sexual orientation, and socioeconomic representation of scientists in taught materials does not match that of the student body [19–21]. These biases persist into undergraduate and postgraduate education. For example, in the UK there are fewer Black, Asian and Ethnic Minority postgraduate students in science than at undergraduate level [22]. This means that the pool of graduate teaching assistants will look less diverse than the undergraduate class. Recommended ‘reading lists’ in science have attracted less attention than in the arts and humanities, but there is some evidence to suggest that science undergraduates are disproportionately directed towards literature from male authors, and to few studies conducted outside of Europe, Australia and North America [21]. Exposure to these repeated biases throughout education reinforce a ‘norm’ that scientists are more likely to be white, male, able bodied and Western.

The lack of diversity in the way that science is traditionally presented means that learners may not have visible role models available to them. As such, learners from historically minoritised groups may feel that they do not ‘belong’ in science, or that scientific careers are not something that they can aspire to [23]. Much of the thinking around scientific role models has centred on binary gender representation, but representation of ethnicity, disability, LGBTQ+ identity is gaining increasing focus [23–26]. Having visible role models from a similar background may increase a student’s sense of ‘science identity’ [27], i.e. a student’s sense of themselves as the “kind of person” who can succeed in STEM [28]. For example, faculty members can act as positive role models for their students. There is evidence to suggest that being taught a traditionally male-biased STEM subject by female instructors increases the likelihood that a female student chooses to continue studying that subject [29,30]. Similarly, having an instructor from a similar ethnic background increases the educational performance of black students [31]. However, if the teaching staff in a given department are not particularly diverse, relying on faculty members as role models will be insufficient, so more proactive efforts need to be made to increase representation.

### Barriers to improving representation in university level STEM education

While the lack of diversity in science is increasingly seen as an issue that needs to be confronted, individual educators often struggle to identify tangible actions they can take to address this within their teaching [32]. There have been many calls to diversify and decolonise science teaching [33], actively confronting the historical legacies of science and the white Western approaches that underpin scientific thinking and conventions [24]. Decolonisation and diversification is an issue for all disciplines, but is often seen as something that is more relevant for arts and humanities than the sciences. Many faculty members actively disagree with diversification efforts within science, seeing science as universal and inherently objective [34]. This mindset fails to acknowledge that science is built on white Western forms of knowledge and thought, and that this bias might be alienating to students of colour [24]. The ‘objectivity’ argument also fails to account for implicit biases about the ‘quality’ of science from Lower and Middle Income countries [35]. Even well-meaning teaching staff in science subjects often feel that they cannot devote time to diversity and inclusivity within the curriculum due to the amount of technical content to be delivered [36]. Given the historical white male bias of scientific research, the need to cover ‘key’ topics in the development of the discipline may hinder efforts to present a greater diversity of scientists. Scientists also often lack the confidence and training to engage with this due to their lack of background in historical, cultural or social science disciplines [37]. Many academics also question whether they are even “allowed” to engage in these discussions when they do not belong to historically marginalised groups, or assume that responsibility for inclusion lies elsewhere in the university [36]. The burden of addressing equality and diversity issues often falls disproportionately to faculty members from historically marginalised groups, creating an unfair burden on individuals who are already structurally disadvantaged by the academy [38]. To avoid this, all members of the academic community have a responsibility to actively address equality and diversity. Individual academics, including those from majority demographic groups, need to feel empowered to take tangible actions, and to recognise that they have an important personal role to play in establishing an inclusive learning environment [32,36].

### Strategies faculty members can used to increase awareness of equality, diversity and inclusion into disciplinary teaching

There are multiple strategies that could be used to embed consideration of equality, diversity and inclusion into university curricula. For example, some instructors are now introducing implicit bias tests such as the Harvard Implicit Association Test (IAT) [39] into the curriculum, and asking students to reflect on their biases and assumptions [40–43]. This strategy has been adopted in several healthcare disciplines, but there are recent reports of similar strategies being used within STEM. One US based chemistry academic describes a positive effect of introducing ‘extra credit’ activities such as taking the Harvard IAT and writing a reflective essay based on the bias highlighted [44]. However, incorporation of these activities takes up time in the curriculum, and may be perceived by students as irrelevant to their scientific training. It is worth advertising at this stage that good resources, with actionable interventions to improve the inclusivity of university teaching are available (see Dewsbury and Brame[45]).

An alternative approach that is more naturally compatible with delivery of science teaching is to introduce more explicit representations of diversity within existing lectures. One advantage of a research-driven curriculum within universities is that teaching materials like presentation slides produced by individual members of teaching staff can change much more dynamically than resources such as textbooks. Case studies of individual scientists from a variety of backgrounds could be included [27], or a diverse range of guest experts invited to contribute to taught sessions [46,47]. These strategies are potentially powerful, but again require space to be found in the curriculum, or rely on academics personally knowing individuals from a range of backgrounds and identities they could invite, so may be difficult to implement. Alternatively, faculty could actively incorporate research from a more diverse authorship into their teaching. There are some efforts to provide resources banks to help academics with this in a subject specific context; for example, Project Biodiversify is an emerging collection of case studies and resources of scientists from a variety of different backgrounds [48]. While this is to be encouraged, academics may still be concerned that incorporation of more diverse authors may result in even more content being added to crowded curricula, or result in less coherent summaries of disciplines if key references are removed to accommodate diversity of authorship [34].

A simpler alternative is to better represent the diversity of authors of papers that are already included within existing lecture materials. For example, a typical powerpoint slide might present a scientific result (e.g. a graph) alongside a formal citation of the work using a standard referencing convention such as Harvard style (Figure 1A). While this format is frequently used, it potentially dehumanises the scientists involved, and obscures any demographic information such as gender (identity), ethnicity, age or other observable characteristics [32]. As such, we hypothesise that students may under-appreciate the diversity of practising scientists when research is presented in this format, instead relying on their implicit biases about the authors. This study therefore considers the impact of including photographs and full names of authors on existing powerpoint slides, giving a ‘humanised’ representation of the scientists. In this format, students are presented with explicit representation of scientists alongside their findings and citation (Figure 1B). This is easy to implement, requiring only relatively minor modifications to existing teaching materials. It does not require time to be spent actively discussing diversity in class, but increases the diversity of representation that students are exposed to while learning about current research. While this method has already been adopted elsewhere[48], to the best of our knowledge, no previous studies have attempted to investigate the impact of the intervention or students’ perceptions of it.

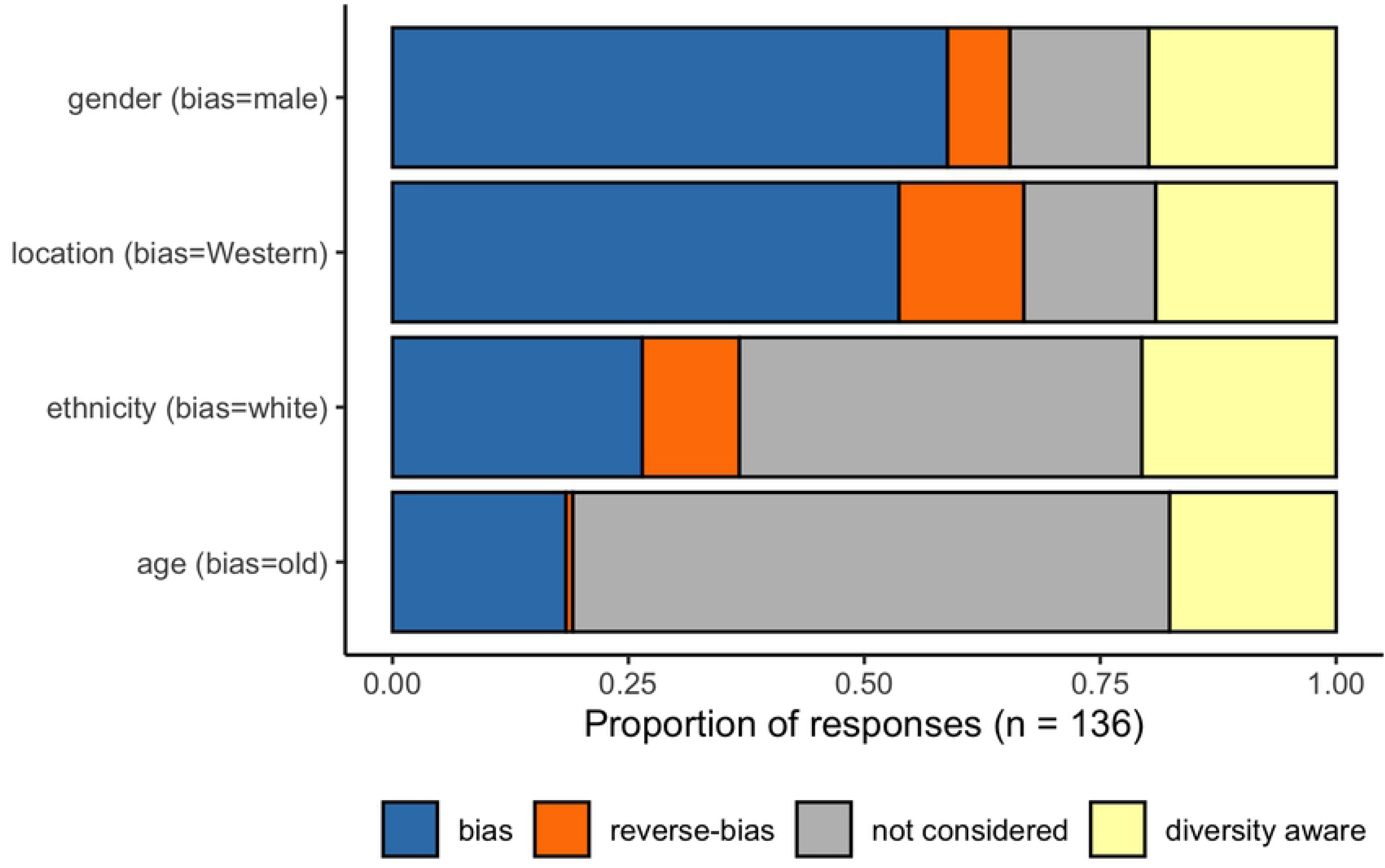

### Aims of Current Study

This study aims to explore the impact of presenting photographs and full names of scientists on existing lecture slides as an educational practice that could be easily adopted more widely. We present the results of a survey completed by 161 undergraduate and taught masters students at a civic university in the United Kingdom. The study has four specific research questions:

1. Do students make inherent assumptions about the identity of scientists based on presentation of author information through formal referencing conventions?
2. Does presentation of photographs and full names of authors of scientific papers change student perceptions of diversity in science?
3. Do students think that presentation of photographs and full names alongside formal referencing conventions is good educational practice?
4. Does the gender or ethnicity of the student have an influence on any of the above?

## Methods

### Ethical oversight

Ethical oversight for this study is provided by the University of Hull Faculty of Science and Engineering Ethics committee (Project code FEC_2019_204). Participation in the study was entirely voluntary, and participants were provided with study information before providing informed consent.

### Questionnaire Distribution

This study was undertaken both before and during the COVID19 pandemic (March 2020 to November 2021). This is relevant because the primary mode of delivery of lecture-based teaching changed during this period of time from ‘face-to-face and didactic’ to ‘online and flipped’. However, the fundamental concept of the survey remained the same as it consisted of a total of eight questions and used a ‘pre’ / ‘post’ format centred around a single taught session on a course. Participation was entirely voluntary with no incentivisation and students were asked to not submit a survey if they had done so previously elsewhere on the course. Prior to the taught session, participants completed the four questions investigating their scientific self-identity, their sense of belonging within the scientific community, and the implicit bias test. After the taught session, participants completed the four questions related to the ‘humanised slide’ intervention. Inclusion of demographic information on gender and ethnicity was an optional extra at the end of the survey. Before the pandemic surveys were paper-based, with the ‘pre’ and ‘post’ questions on different sides of the same sheet. During the pandemic surveys were presented as two separate Canvas (Virtual Learning Environment) quizzes separating out the ‘pre’ and ‘post’ questions.

### Survey questions

Prior to the beginning of a taught session, participants were asked to share their first impressions on the identity of the author of a hypothetical reference in Harvard format; Lee, M. (2019) ‘Globally interesting Biology’. *Biology Journal*: 45 p. 12-25. ‘M. Lee’ was chosen specifically to minimise inherent assumptions about the author as the surname Lee is common in communities descended from Anglophone, Korean, and Chinese ethnicities. There are many first names beginning with ‘M’ through most ethnicities with limited inherent bias as to the gender of the name. Participants were asked specifically to reflect on “what the author looks like, their first name, and where they come from?”. To the best of our knowledge, no other study has undertaken an investigation similar to this and therefore there may be uncertainties about using an unvalidated tool. However, as much as possible the question was designed to not lead the participants to any particular response. In fact, in the results section many responses did make explicit assumptions about the author but a large proportion of participants highlighted that one cannot make assumptions based on the information provided. The survey followed a straightforward approach to gaining student perspectives on the humanised slide intervention. Participants were asked whether the intervention had changed their perspective on their answers to the set of questions in the ‘pre’ survey. They were then asked whether they felt that having ‘humanised’ slides could be considered ‘good practice’. All survey questions can be found in full in Supplementary Table A.

### Thematic coding

To analyse the free text data qualitatively, we undertook a thematic analysis based on the protocols outlined by Braun and Clarke [49]. Thematic analysis is a widely used, flexible and rigorous approach to analysing qualitative data through the development of themes and subthemes within a dataset [49,50]. Every understandable response for each of the open text questions was coded to one of three or four themes only, responses to these questions could not be coded twice to different themes as they were clearly mutually exclusive. For example, responses to the inherent bias question were categorised as either: bias, reverse-bias, not considered, or diversity-aware. Allocation of a response to a theme was highly unambiguous, and the exact terms that define each theme are outlined in Supplement A. The mutual exclusivity of the coding themes means that the data meet the assumptions for a chi-square test of independence, with the theme of the response being a categorical dependent variable, and gender/ethnicity being categorical independent variables [51]. For each question, two chi-squared tests were performed to determine whether the proportions of responses coded to each of the four themes were altered by participant Gender or Ethnicity (one test for each categorical variable). Bonferroni corrections were used to account for the use of multiple tests per question. Note that not all participants provided demographic information, and responses that left this blank or responded prefer not to say were not included in these analyses. Due to limited representation in the data set, Gender and Ethnicity variables were aggregated into two categories centred around an analysis of the concept of white, male fragility [52]; however, even with this correction there was insufficient representation within the BAME group for statistical tests investigating ethnicity as a predictor variable. Gender was represented as either ‘Male’ or ‘Female and Non-binary’. Ethnicity was separated into ‘White’ or ‘Black, Asian, or Mixed Ethnicity’ [i.e. BAME].

## Results

Students on eight different taught modules where the ‘Humanised slides’ were used were invited to complete the survey. 161 students responded to the survey, including 111 undergraduates and 50 students on taught masters courses (Table 1). This represents a 41% response rate, so findings are not necessarily reflective of the whole cohort. 125 participants provided their gender identity, and 124 provided their ethnicity (Table 2). Within that subset, the sample was biassed towards female (58%) and white (87%) participants. As there were relatively few responses from Black, Asian and Mixed Ethnicity students these have been grouped together for analysis, however we recognise that this may obscure differences between cultural groups. We also grouped together female and non-binary students for analysis, but excluded those who preferred not to disclose their gender identity.

**Table 1:**
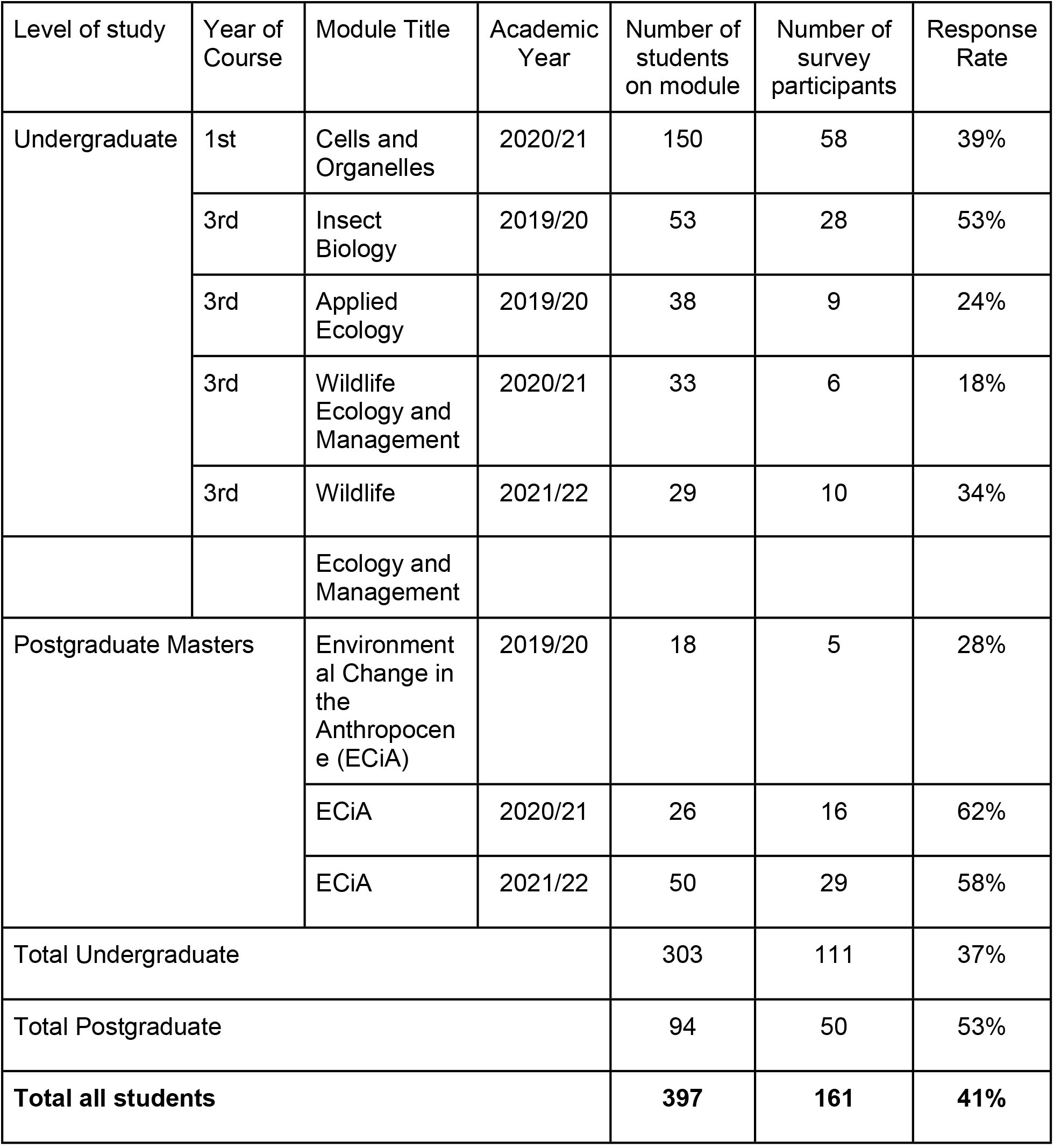
Survey population and response rates

**Table 2:** Participant demographics

| Demographic | Category | Number of responses | % of total responses | % of those that provided demographic information |
| --- | --- | --- | --- | --- |
| Gender | Female | 73 | 45% | 58% |
|  | Male | 48 | 30% | 38% |
|  | Non-binary | 2 | 1% | 2% |
|  | Prefer not to say | 2 | 1% | 2% |
|  | Not provided | 36 | 22% | - |
| Ethnicity | White | 108 | 67% | 87% |
|  | Black, Asian or Mixed Ethnicity | 16 | 10% | 13% |
|  | Not provided | 37 | 23% | - |
| Total |  | 161 | - | - |

### Student Implicit Assumptions of Author Identity

We first wanted to determine if students held implicit biases around author identity when presented with a conventional academic reference. There were 136 understandable and complete responses to the survey question asking students to share their first impressions on the identity of the author of a hypothetical reference in Harvard format; Lee, M. (2019) ‘Globally interesting Biology’. *Biology Journal*: 45 p. 12-25. All responses were coded to one of four categories for each of four different characteristics. The categories were bias, reverse bias, not considered, and diversity assumed. The characteristics were gender (bias = male), geographic location (bias = Western-ised), ethnicity (bias = white), and age (bias = old/middle-aged).

“*White male, middle aged, Mark??? England or America*” - an example of a response coded as ‘Bias’ for all four characteristics.

The most common assumptions students made about author identity related to gender and location (Figure 2). Of the 136 responses, 59% explicitly assumed that the author was male, and 54% assumed they were from a Western-ised country (i.e. USA, Europe or Australia). Participants were less likely to make explicit statements about the ethnicity of the author, but the most common explicit assumption was that the author was ‘white’ (26% of all respondents). White ethnicity was only coded when the terms ‘white’ or ‘caucasian’ were used and not based on location assumptions, so the difference in location and ethnicity data may be due to a lack of clarity in the format of communication. Author age was not asked for, but many participants made references to the age of the author, as well as regular references to facial hair. Age was the least frequently mentioned aspect of author identity, but the most common assumption was that the author was old or middle-aged (18%).

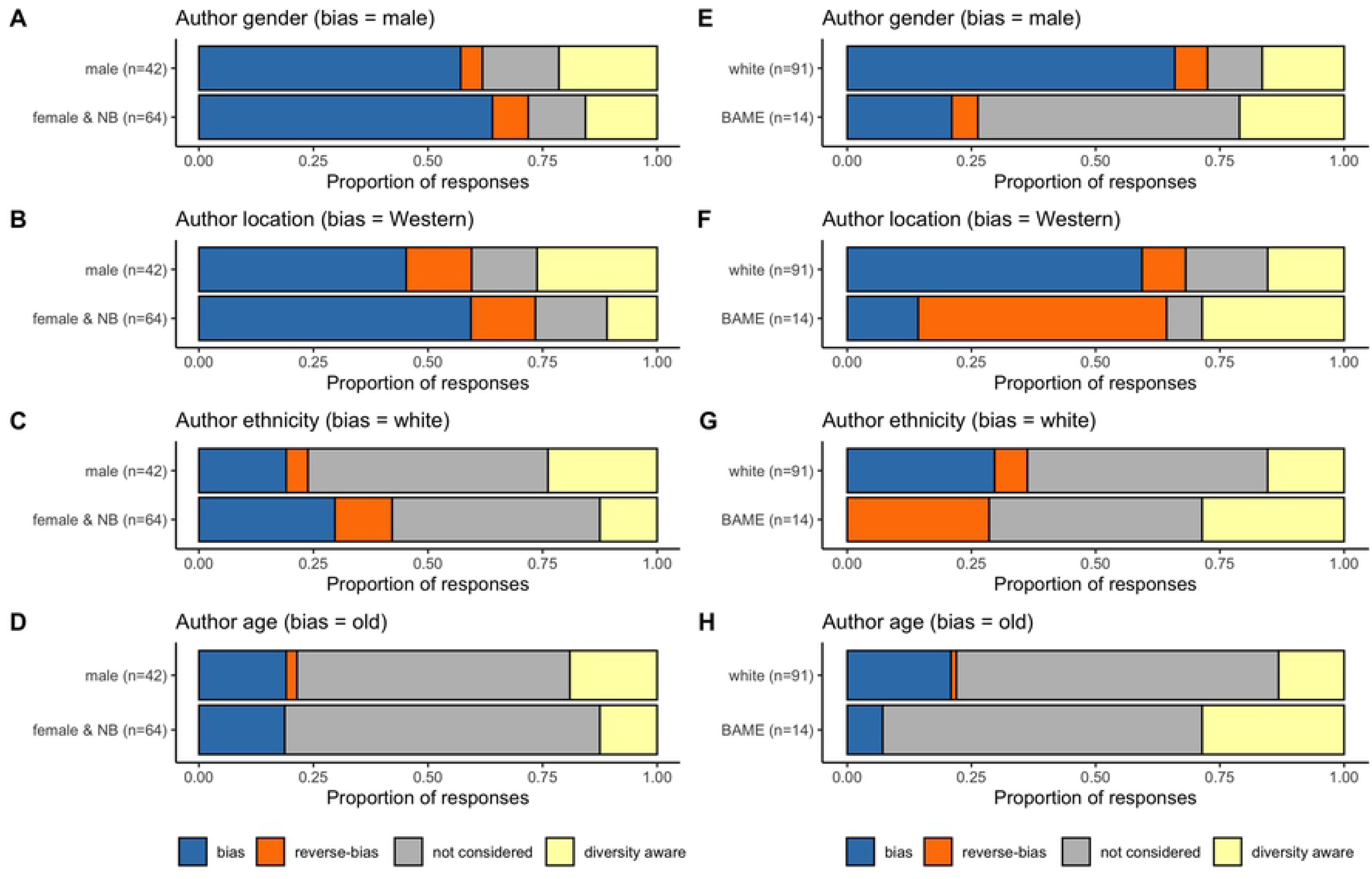

Reverse-bias responses were most common for location 13% (i.e. not Western), then ethnicity 10% (i.e. not White), then gender 7% (i.e. not male), and one respondent explicitly stated the author was young (i.e. not middle-aged or old). Respondents who did not make explicit assumptions either said that they were unsure about author identity, or that one could not make assumptions about an author from a Harvard reference; which we coded as ‘Diversity aware’. Diversity aware responses were given by approximately 20% of participants (age 18%, location 19%, gender 20%, and ethnicity 21%). However, within diversity aware responses some participants still made implicit assumptions about the identity of the author. For example, one response stated that you cannot make assumptions about ethnicity or location, but then uses masculine pronouns demonstrating implicit gender bias; “*There isn’t a first instinct, name can’t be used to distinguish what someone looks like or where they come from. **He’s** just as likely to be a white american as **he** is to be a black african*”.

Within the responses that did make assumptions about author identity, the majority of responses represented the bias response. The strongest bias was for age (96%biased: 4% reverse-biased), followed by gender (90%:10%), location (80%:20%) and ethnicity (72%: 28%).

We were interested to see if the personal characteristics of the students made any difference in their implicit assumptions about author identity. We therefore broke down the responses to the assumptions data by participant gender and ethnicity (Figure 3). There was insufficient data to reliably perform statistical analysis of responses by participant gender or ethnicity, as for all comparisons there were multiple categories with fewer than 5 respondents making a Chi-square test inappropriate. However, it should be noted that none of our BAME participants assumed that the author was white (Figure 3G), and the majority of BAME participants gave the reverse biased response for location (assuming a non-Western location; Figure 3F). Our data therefore indicates that a majority of students from all backgrounds do make implicit assumptions about who is participating in science from Harvard style references, but that these assumptions might differ between students from different demographic groups.

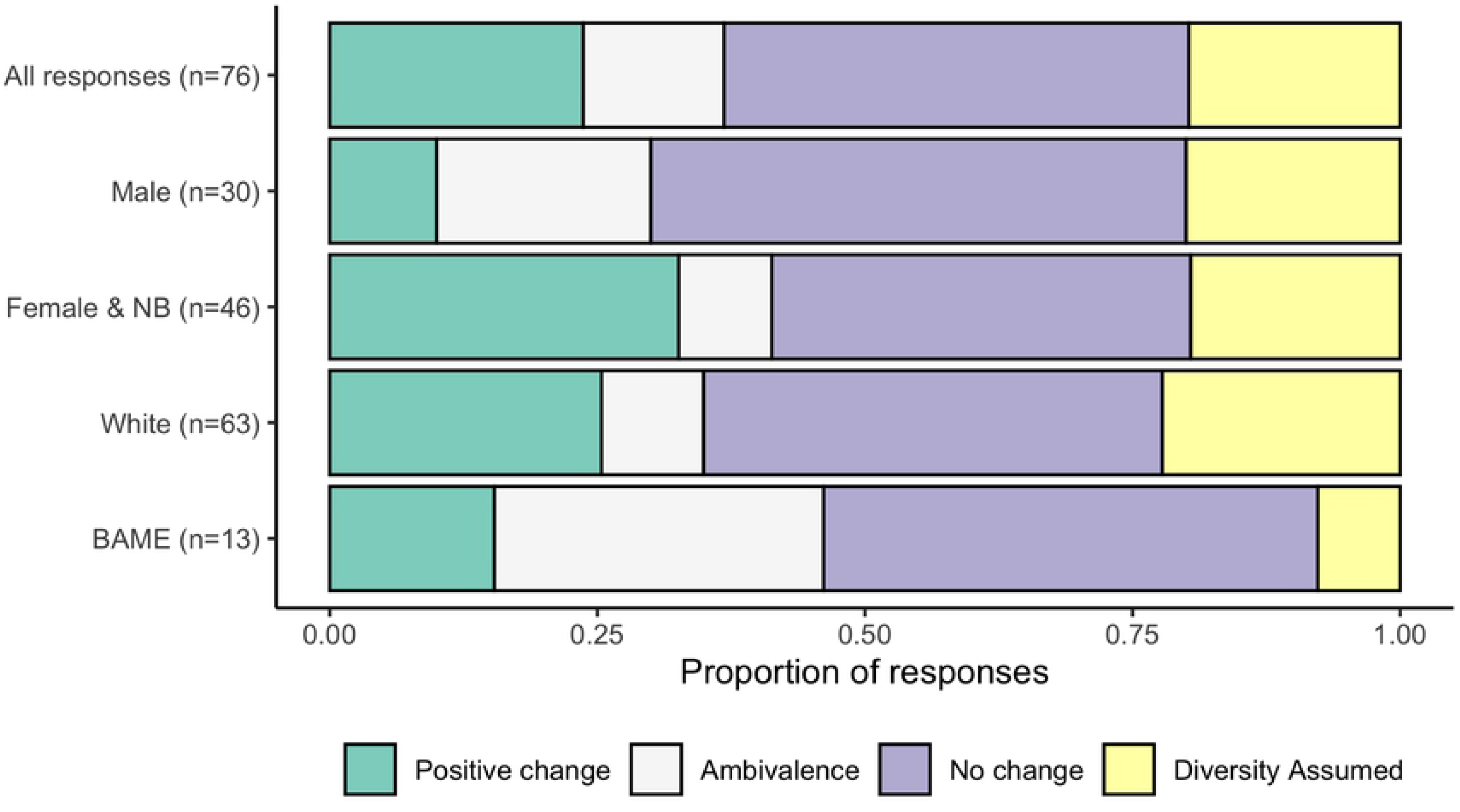

### Impact on participants perception of diversity in biosciences

Having established that students did make implicit assumptions about author identity from Harvard style references, we then investigated the impact of using the ‘humanised’ slide design in the lecture. We asked students whether seeing explicit representations of scientists changed their perceptions of diversity within science (Figure 4). There were 76 coherent responses which were all coded as either ‘Explicit no change’, ‘Explicit positive change’, ‘Diversity assumed’, and ‘Ambivalence’. The most commonly coded response was ‘Explicit no change’ [33/76], mostly without further justification (i.e “*No*”); although some responses outlined that the participant felt the practice was irrelevant and/or unnecessary [3/33]. Of the 33, ‘Explicit no change’ responses, five suggested that the practice was insufficient or did not display ‘enough’ diversity. Interestingly within this category, two participants’ responses were very focused on gender as diversity and less receptive to ethnicity/location as diversity (e.g. “*No, majority were still male* (*although more diverse*) -> *mainly asain [sic]*”). Note that the claim that the “majority were male” is inaccurate; as a minimum all lectures were designed to present at least 50% of the scientists as female.

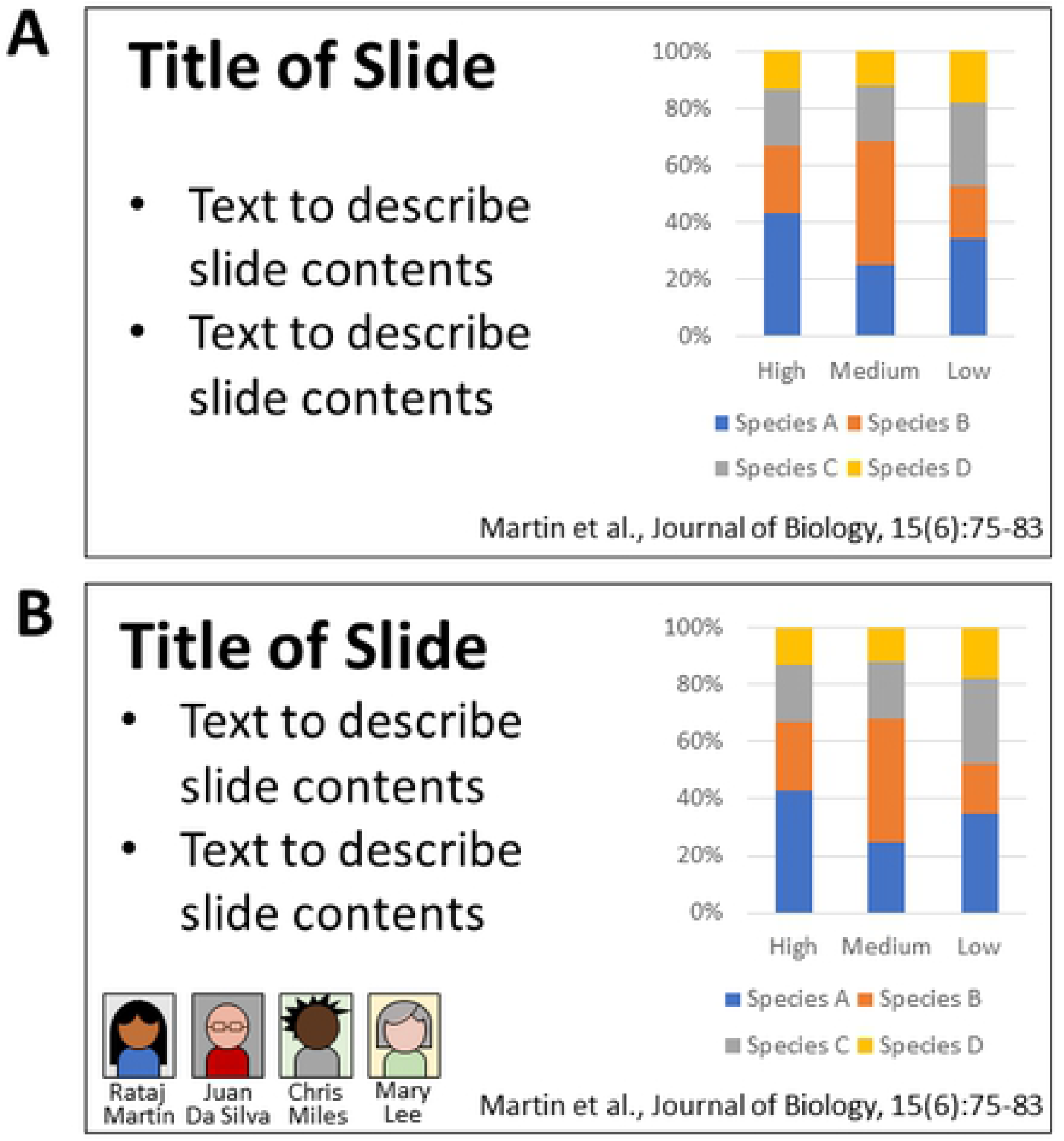

The second most commonly coded response was ‘Explicit positive change’ [18/76] where participants stated that their perceptions of diversity in the field was improved; most often this was gender-related but students also commented on how their perceptions of ethnicity, gender-identity, and/or age were impacted (e.g.“*Yes - it highlights the presence of ‘non-cis white males’ also publishing papers.*”). A common sub-theme within the ‘Explicit positive change’ theme, was participants expressing the belief that there is no standard look for a scientist or that anyone can be a scientist [6/18] (e.g. “*There isn’t a certain “look” for a biologist - anyone can be one.”)*.

The final theme that was expressed consistently throughout the responses was the idea that the practice did not change the participant’ sperceptions because they already knew that the field was full of diversity (diversity assumed) [15/76] (e.g. “*Not at all, anyone can be a biologist like anyone can cook*”). The data set is not sufficiently large to determine whether respondents coded in this theme were also coded as ‘diversity aware’ in response to the implicit bias question. However, some participants made explicit/implicit assumptions about the identity of the hypothetical author earlier in the survey and then said that one could not make assumptions (e.g. “*First name: Michael. From: Europe.*” followed by “*No, I already had the mindset that it would be a wide range of people carrying out the research.”)*. The remaining responses were either attempts at humour, expressing a lack of understanding of the question, or expressing disinterest in the survey, which were coded as ‘Ambivalence’. There was no difference in the proportion of responses coded in each of the four categories according to participant gender (Figure Y; X^2^_1(N=76)_=6.17, p=0.10). The low number of responses from BAME students meant that statistical comparison is inappropriate, but the data is presented in Figure 4

### Student perceptions of the practice

We investigated whether students felt that there was value in explicit representations of diversity in lectures as an educational practice by asking whether they felt it was ‘good practice’. There were 91 coherent responses which were all coded as either ‘Explicit good practice’, ‘Ambivalence’, or ‘Explicit not good practice’.

The most commonly coded response expressed explicit support for the practice (68/91 = 74.7%). While 18 of these responses were positive with no additional details (e.g. “*Yes*”), the vast majority of respondents made some attempt to explain their decision. This elaboration was further broken down into a number of sub-themes evident in Table 3; note that responses could be coded in multiple sub-themes but not in multiple themes. ‘Explicit good practice’ responses were most commonly qualified with statements expressing the value of the practice for raising ‘awareness of diversity’ in science/bioscience (34/68= 50%). There was also a related but subtly different theme expressing that the practice ‘humanized’ the researchers and/or made them more personally relatable to the participants (19/68 = 28%). Some ‘humanized’ responses focused on how the practice might help other people, while for others it had a more personal impact on how they felt they fit within STEM.

**Table 3:** Frequencies of sub themes within the explicit support for practice, presented by participant gender.

| Sub theme | Illustrative examples | Male<br>(n=25) | Female<br>+ Non-Binary<br>(n=41) | Total<br>(n=68) |
| --- | --- | --- | --- | --- |
| 'Awareness<br>of diversity' | *"Yes, breaks down stigma that you have to fit a specific mold to be successful in science"*<br>*"Absolutely - yes. Using pictures and full names 'normalises' the huge variety of people who contribute science to society. This provides a platform for equality and inclusivity and makes people think they can do science too - which of course they can."* | 7<br>[33%] | 27<br>[60%] | 34<br>[50%] |
| 'Humanising<br>researchers' | *"Seeing a face and a name behind the research, makes it less intimidating, and potentially more reader friendly"*<br>*"Seeing just one visibly queer person would help me i think. nothing against old white men but it would help me feel safe"* | 5<br>[24%] | 14<br>[31%] | 19<br>[28%] |
| 'Credit +<br>recognition' | *"Yes, they work hard and deserve to be recognised"*<br>*"Yes, it give's more perspective over when their research was carried out and makes me feel more invested in their research as oppose to one of thousands of relatively anonymous scientists as they aren't appraised enough for their contribution"* | 7<br>[33%] | 2<br>[4%] | 9<br>[13%] |
| 'Utilitarian' | *"Yes, gives us more knowledge for future work we may have to look into that could be linked"*<br>*"Yes, can tell exactly who wrote the article and easier to find it for own reading."* | 5<br>[24%] | 7<br>[16%] | 12<br>[18%] |

There were some additional and unexpected reasons why respondents felt the practice was positive that were unrelated to diversity in STEM (Table 3). The idea of credit, praise or recognition for the scientists undertaking the work was a recurring sub-theme, focused on the idea that scientists deserved less anonymity in their contribution to the field (9/68 = 13%). The final sub-theme expressed that having full names and headshots might help them remember the content of the papers better or find more research by the same authors, which was coded as ‘utilitarian’ (12/68 = 18%). Initial investigation of these sub-themes by participant gender suggests that male participants were more likely to focus on credit & recognition than female and non-binary students (33% of male responses, 4% of female/non-binary). Conversely, female and non-binary participants were more likely to provide responses aligning to the sub-themes ‘awareness of diversity’ (60% of female/non-binary, 33% of male) and ‘humanising researchers’ (31% of female/non-binary, 24% of male). Note that while a gender bias is suggested in the data, because each response could be categorised in multiple sub-themes statistical analyses are inappropriate.

A much smaller number of respondents disagreed that this was good practice [14/91], coded as Explicit not good practice. Justifications sometimes included that the practice was unnecessary or detracted from the science (e.g. “*I would rather focus on the information rather than the author*”). A further nine respondents were unsure, ambivalent, or had mixed feelings about the practice (e.g. “*Interesting but unsure of relevance*”). A deeper investigation of student responses to this question according to the self-identified gender of the respondent suggests that males were significantly more likely to express negative or ambivalent feelings about the practice than female or non-binary students (Figure 5; X^2^_1(N=88)_=5.18, p=0.02). Statistical comparison by student ethnicity was not possible due to low sample size.

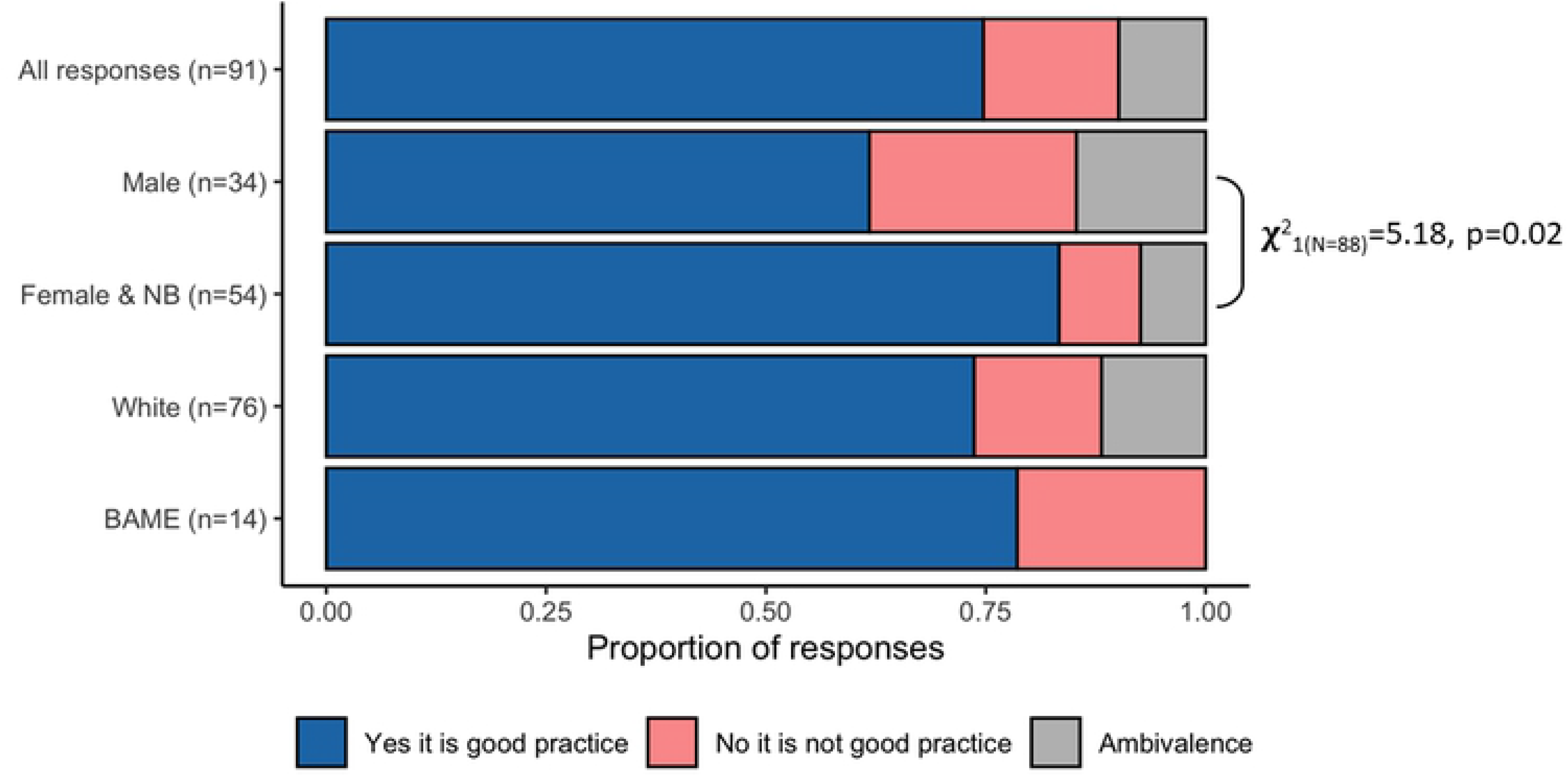

## Discussion

This study investigated the value of a simple intervention attempting to raise the visibility of diversity in STEM by explicitly including the full names and photographs of the authors of research used in formal teaching materials; i.e. the use of ‘humanised’ slides (Figure 1). The intervention stems from an initial assumption that many formal referencing systems exacerbate prevailing perceptions that STEM is not diverse, which is unconsciously embedded into students throughout their education [19–21]. The first part of this study investigated whether students made inherent assumptions about the identity of researchers based on author information presented in formal referencing conventions alone. We found that most participants made some kind of assumption about the gender, ethnicity, location, or age of the paper author cited in the Harvard style; with clear biases towards the author being male, Western, white, and middle-aged or older. These assumptions about who ‘researchers’ are match those present in validated measures of implicit diversity biases in STEM, such as ‘DAST’ [13–15] and the Harvard IAT [39]. Our results also align with the predictions of other studies looking at the diversity of STEM researchers in the taught environment like reading lists and textbooks [16]. It is worth noting that in our study white, male, and Western biases appear to be consistent across participant genders, and that even students who were consciously aware of the potential for implicit bias still made assumptions about the author’s identity. Combined these results suggest that student implicit biases about the diversity of STEM are exacerbated by the absence of explicit diversity cues in ‘neutral’ referencing systems.

Having established that diversity-neutral researcher information encourages biassed perceptions of STEM diversity, we then asked whether our visible diversity intervention was able to challenge student implicit biases. When asked whether the ‘humanised’ slides impacted participant perceptions of STEM diversity, over 60% of the participants felt that their perceptions were unchanged. Some felt that the intervention was insufficient to engender change, while others that they had already assumed that STEM was a diverse field. However, for an important subset of those surveyed (over a fifth) the intervention was impactful, with many highlighting that it challenged stereotypes about who can be a scientist. The impact was not necessarily just felt among students within groups that are seen as under-represented within STEM, as we did not find a significant difference in impacts between male, and female or non-binary students. Our sample was too small to formally compare the impact along ethnicity lines, but this is an area that could be investigated further with a larger study.

Some of the key barriers to engagement with diversity enhancing interventions are student perceptions of them and their willingness to engage with them; particularly those students within majority groups who may feel ‘threatened’ [53,54]. However, when asked in this study, 75% of participants agreed that the use of ‘humanised slides’ was good practice. Most commonly this was linked to the potential for the intervention to raise awareness of diversity in STEM or supporting an individual’s sense of belonging within the field, which would be expected from studies of the use of pictures to portray actual diversity in STEM [55]. In addition to those reasons expected, some students highlighted the value of the practice for providing proper recognition to hardworking researchers or for improving their ability to investigate the research more easily. There was a statistically significant gender split in the data, which suggests that male participant perceptions were more likely to be negative or ambivalent than female and non-binary participants. While the overall proportion was small, this fits with the wider literature on the concepts of white, male fragility [56]. Negative responses particularly focused on the intervention as being unnecessary or irrelevant, which is well-established within the literature on barriers to anti-racist and anti-sexist pedagogies [54,57]. This response likely stems from initiatives that highlight white, male privilege are interpreted as personal attacks or as minimising the struggle and effort they have invested into their own successes [56]. Interestingly, it appears that the male participants that felt it was good practice appeared more likely to focus on credit/recognition as a benefit of humanised slides while female and non-binary students were more likely to raise diversity and/or belonging values. This may be why we found limited resistance to the intervention among male students, because there was an explicit benefit that did not clash with their self-identities.

### Limitations of study

This is intended as an exploratory study to explore whether students hold implicit biases about the authors of scientific papers, and practices which might mitigate against that bias. While our conclusions and methodology are robust within the scope of the study, our findings are not necessarily applicable in all contexts. The sample size is modest, but compares favourably with some other published reports of interventions to improve student understanding of diversity in university STEM settings [44,58]. While our sample is drawn from postgraduate and undergraduate students at multiple stages of their degrees, participants are all at a single UK university. Participation in this study was voluntary, resulting in a response rate around 41%, so responses may not reflect those of the wider cohort. Our data collection also happens entirely within the context of biosciences and ecology which are typically more gender balanced, so may not represent other STEM disciplines. It would be particularly informative to extend this study across the STEM disciplines, including those with a more significant gender and ethnicity bias. For example, the Geosciences have been highlighted as being particularly biassed towards white researchers [59], and the significant male bias in physical sciences, engineering and mathematics is well known [1]. The Harvard style reference implicit associations component of this research is a novel research methodology which should be developed and validated further, but the white western male bias in responses is consistent with that exposed by other well established methodologies such as the Draw A Scientist Test and Harvard Implicit Association Test [14,15,39]. It should be noted that our study captures immediate responses to the impact of the ‘humanised’ powerpoint slide design, but does not attempt to measure a longer term impact or changes in student opinions about whether they are likely to undertake a scientific career. However, we consider the research to have value as a ‘snap shot’ study that highlights potential opportunities to increase diversity and representation within STEM education, as well as presenting an opportunity to have more open discussions with students about diversity and decolonisation of the curriculum.

### Practical considerations of using the humanised slides as an educational practice

Although our data indicate that the humanised slide design has a positive impact on student perceptions of diversity in science, we recognise that this intervention is insufficient to address the systematic biases within STEM. Our data suggests that presentation of humanised powerpoint slides in a single undergraduate lecture has some positive impact on student perceptions of diversity, and therefore who ‘belongs’ in the scientific community. It is important to note that we do not see the relatively modest number of students for whom the intervention was impactful as an indictment of its efficacy. Even if only one student’s perception of their own place within the field or their implicit biases about who belongs in STEM changes as a result of this intervention, it will have had a meaningful impact. However, this intervention should be seen as a very small part in a wider catalogue of interventions that can be used to raise visibility of diversity in STEM [27,45,48]. The use of ‘humanised slides’ is designed as a simple pedagogy that could be implemented by anyone efficiently. We see it as an important precursor to the long-term, and much slower, efforts to raise STEM diversity in textbooks, faculty membership, and wider portrayals of scientists in education and the media. There is also a danger that isolated attempts to improve diversity are seen as irrelevant or tokenistic, as it is known that poorly designed mandatory EDI interventions are ineffective or can even backfire and create resentment or workplace tension [60]. For this practice to have sustained positive change for the majority of students it needs to be used repeatedly and consistently through a programme by multiple teaching staff. It must also be accompanied by other active efforts to improve diversity and representation including hiring practices, inclusive curriculum design, appropriate mentoring schemes and decolonization of the curriculum. A genuinely inclusive STEM curriculum would make space in to actively discuss equality and diversity from a variety of perspectives going beyond gender and ethnicity to include non-binary and trans, disability, socioecoomic class and international viewpoints [48]. Relying entirely on the humanised slide design is insufficient, as students may still make implicit assumptions from photographs and names, and photographs cannot capture hidden aspects of diversity such as sexual orientation or non-visible disabilities. This approach is not the only way to increase representation, and will be complemented by a range of other strategies. We would also encourage the inclusion of alumni or expert lecturers from diverse backgrounds in teaching, as well as ‘scientist spotlight’ assignments [27] and other curriculum efforts to improve diversity [48].

It should also be noted that the humanised slide design practice is also only effective if instructors actively reflect on the studies they are including and make positive efforts to improve representation within their teaching materials. If the only studies presented in this format are written by older white men, then this practice could reinforce a perspective that practising scientists do not come from a diverse range of backgrounds. As academics, we have found that preparing slides in this format has prompted us to reconsider the studies we include in lectures, and actively seek out papers with a more diverse authorship. As such, we feel that the practice is also of benefit to instructors who are trying to make a positive difference but feel unsure of where to start. It should be noted that while designing slides in this format is a relatively modest intervention, it does increase the time spent on preparing teaching materials. We have found that selecting literature with appropriate authorship and obtaining images of researchers through searches on GoogleScholar or institutional websites does take time, but consider the activity to be of sufficient benefit. We recommend this as a straightforward way that academics in any setting, including those from majority demographic groups, can improve representation in a discipline relevant format.

## Conclusions and recommendations for practice

In this study, we conclude that students make implicit assumptions about diversity of scientists from Harvard style references, and that presentation of photographs and full names of scientists alongside formal citations can have a positive impact on some students. We recommend adopting a ‘humanised’ slide design in research-led teaching materials, so that students can have a better appreciation of the actual diversity of practising scientists that is masked by the formal citation. In our experience, adopting this slide design also actively encourages instructors to seek out papers with a diverse authorship. This is a straightforward intervention that all academics could make to give better representation to a diversity of scientists without having to find additional space within the curriculum. While this intervention is not sufficient to address all issues around diversity in STEM, it is an easy strategy that improves representation in an authentic way through research-led teaching, and may ultimately have a positive impact on the diversity of students choosing a scientific career.

## Supplement A - Survey and coding methods

**Supplemental Table A.**
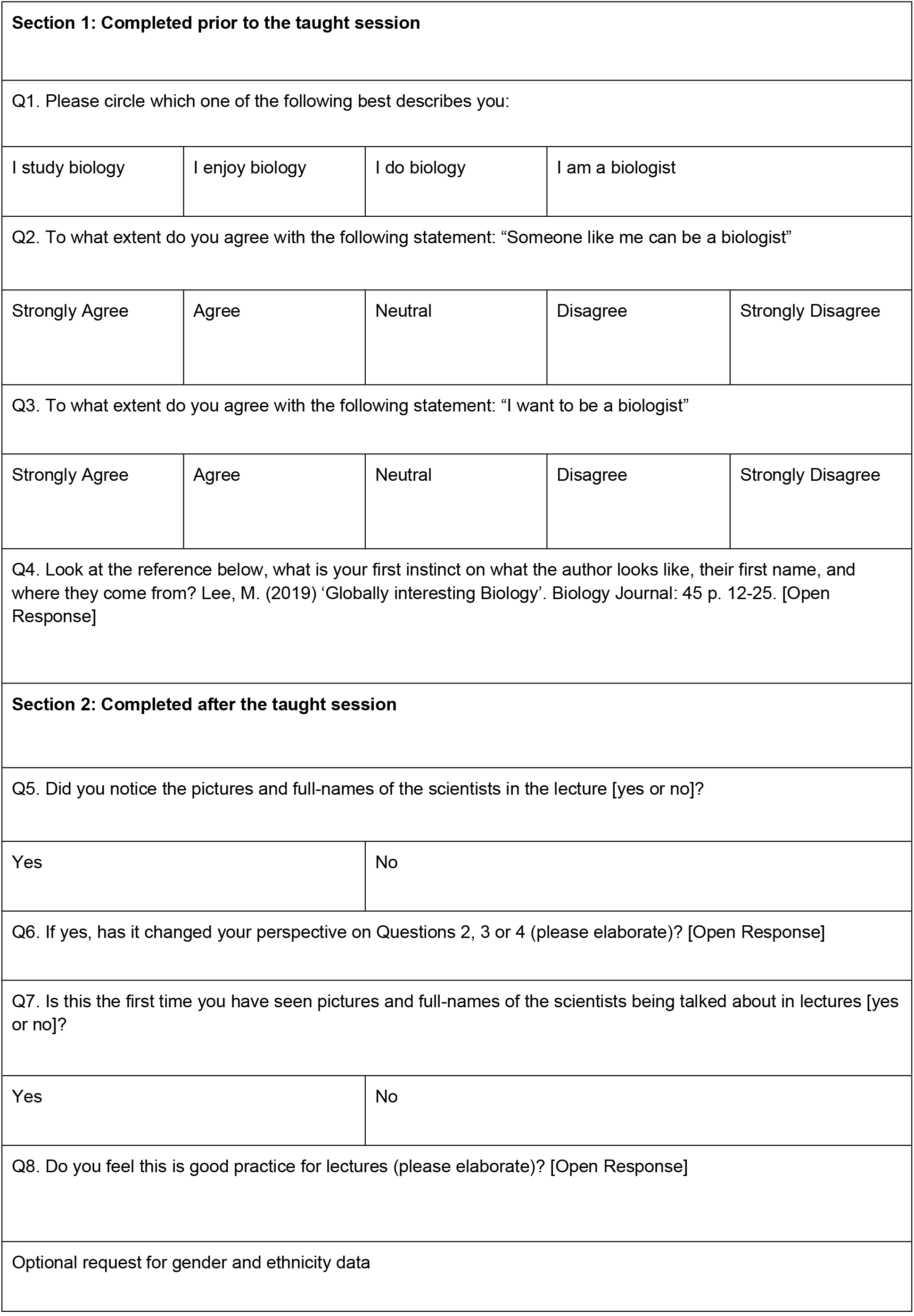
Survey questions and answer format

|  |  |  |  |  |
| --- | --- | --- | --- | --- |
| **Section 1: Completed prior to the taught session** |  |  |  |  |
| Q1. Please circle which one of the following best describes you: |  |  |  |  |
| I study biology | I enjoy biology | I do biology | I am a biologist |  |
| Q2. To what extent do you agree with the following statement: "Someone like me can be a biologist" |  |  |  |  |
| Strongly Agree | Agree | Neutral | Disagree | Strongly Disagree |
| Q3. To what extent do you agree with the following statement: "I want to be a biologist" |  |  |  |  |
| Strongly Agree | Agree | Neutral | Disagree | Strongly Disagree |
| Q4. Look at the reference below, what is your first instinct on what the author looks like, their first name, and where they come from? Lee, M. (2019) 'Globally interesting Biology'. Biology Journal: 45 p. 12-25. [Open Response] |  |  |  |  |
| **Section 2: Completed after the taught session** |  |  |  |  |
| Q5. Did you notice the pictures and full-names of the scientists in the lecture [yes or no]? |  |  |  |  |
| Yes |  | No |  |  |
| Q6. If yes, has it changed your perspective on Questions 2, 3 or 4 (please elaborate)? [Open Response] |  |  |  |  |
| Q7. Is this the first time you have seen pictures and full-names of the scientists being talked about in lectures [yes or no]? |  |  |  |  |

|  |  |
| --- | --- |
| Yes | No |
| Q8. Do you feel this is good practice for lectures (please elaborate)? [Open Response] |  |
| Optional request for gender and ethnicity data |  |

### Survey development

The survey used a pre- / post- model surrounding each taught session.

In the ‘Pre’ part of the survey, participants were asked a series of questions designed to investigate their ‘science identity’ and our implicit bias test (designed around a hypothetical Harvard reference). Our implicit bias test asked participants to share their first impressions on the identity of the author of Lee, M. (2019) ‘Globally interesting Biology’. *Biology Journal*: 45 p. 12-25. ‘M. Lee’ was chosen specifically to minimise inherent assumptions about the author as the surname Lee is common in communities descended from Anglophone, Korean, and Chinese ethnicities. There are many first names beginning with ‘M.’ through most ethnicities with limited inherent bias as to the gender of the name. It is worth noting that in the responses included ethnicities found in both Western and Eastern hemisphere, and both male and female first names were suggested.

The ‘Post’ aspect of the survey was only available after the taught session and first asked whether the participants noticed “the pictures and full-names of the scientists in the lecture materials?”. If they answered yes, they were then asked whether the practice had changed their perspective on the questions asked before the taught session began. While this did include the questions designed to investigate participants’ sense of belonging (Q.1-3), participants’ responses were overwhelmingly focused on Q4. The survey questions related to student perception of the practice were similarly blunt and unequivocable. We asked “Do you feel this is good practice for lectures (please elaborate)?”; after reminding students what the practice was exactly in Q7. In this instance, we were specifically interested in student perceptions of the intervention because perceptions are one of the most significant barriers to engagement with diversity-positive educational interventions [54]. Note that we did not ask the question specifically in the context of diversity-positive education, and participants were free to outline positive and negative responses entirely unrelated to the visibility of diversity in STEM.

### Thematic analysis of responses

This report focuses on participant responses to the three questions on the survey that had open text responses, which relate to research questions 1-3 respectively. The thematic coding process was completed separately for each question, and each will be outlined in detail below. Importantly, to allow for quantitative comparisons a system whereby each response could only be coded into a single theme was developed. This design was adopted to align to assumptions of statistical tests for count-based categorical variables (e.g. Chi-square test).

The first question to undergo thematic analysis was the implicit bias test based on student perceptions of the author of a hypothetical Harvard reference. The process started with an initial review of the participant responses to collate the most common kinds of assumption about the author. The four most consistently assumed characteristics were gender, geographic location, ethnicity, and age; which became the four primary coding themes. Then every response was reviewed for comments relating to these four characteristics and was coded into one of: Bias, Reverse-Bias, Not Considered, and Diversity Aware. A ‘Not Considered’ code meant that the participant had made assumptions about other characteristics but not the specific one being coded (e.g. made assumptions about age but not ethnicity, so was scored as ‘not considered’ for ethnicity). ‘Diversity Aware’ answers either stated explicitly that one could not make assumptions about the author, were sincerely unsure of what to say, or made a humorous comment that made absolutely no assumptions about the author. With respect to bias and reverse bias codes:

- Gender identity was coded either from a direct statement of gender (e.g. male = Bias), the use of gender-specific pronouns (e.g. she/hers = Reverse Bias), or according to the proposal of an explicitly and obviously gender-specific name (e.g. Mike, Martin, etc.).
- For ethnicity, responses linked to ‘white’ or caucasian were coded to Bias, while any other ethnicity (most commonly Chinese) was coded to Reverse-Bias.
- For location, explicit statements of West / Western / Developed were code to Bias, as were any European, UK, or North American locations. The few Australia responses were also included in the Bias theme. Any other location was considered to be Reverse Bias (most commonly China)
- Geographic location and ethnicity were separated by the statement of a place compared to the use of an ethnicity (e.g. China is a ‘Reverse Bias’ location vs. Chinese is a ‘Reverse Bias’ ethnicity). Ethnicity was not assumed from location and vice versa (e.g. it was not assumed that the participant thought the author was Chinese ethnicity if they stated the location as China).
- Biased age was coded from explicit statements that the author was middle-aged, old, bald, or grey-haired, while reverse-bias was coded in the one instance a participant assumed the author was ‘young’.

The second question responses for thematic analysis was Question 6 - “has it [humanised slides] changed your perspective on Questions 2, 3 or 4 (please elaborate)?”. The format of the question meant that the majority of participants started their response with a clear Yes or No answer, which was used as the initial basis for coding. ‘Yes’ responses were coded as ‘Explicit Change’; there were no ‘Yes’ responses that suggested that perceptions of diversity had become ‘worse’ as a result of the intervention. ‘No’ responses were placed into the ‘Explicit no change’ theme, unless the participant had elaborated to explain that the ‘No’ response was because they had assumed that the field was diverse already; coded as ‘Diversity Assumed’. However, not all responses had a clear ‘Yes’ or ‘No’ preface. So responses that included any explicit expression of a positive impact of the intervention on their perception of diversity or their sense of belonging within the field were coded as ‘Explicit Change’ (e.g. “*It’s changed my perspective on 4 because I assumed it would be a white man but they were super diverse*”).Conversely, any response that highlighted a negative perception of the intervention, such as it being irrelevant or unnecessary, was coded as ‘Explicit no change’. The final code allocated was Ambivalent, which was linked to responses that highlighted both positive and negative elements, anyone who was unsure of the impact, or a response that was unrelated to the question. Once the highest level themes were coded each was reviewed separately for similarities in the rationale for participant answers. These sub-themes tied together the different types of reason the participants had given for answering the way that they had. Responses could be coded into multiple sub-themes, particularly where the open text response was long and comprehensive. The sub-themes that arose are considered in detail in the results section.

**Supplementary Table B.** Examples of responses to a question asking whether the intervention had changed student perspectives to the ‘pre-’ taught session questions, focusing particularly on how responses were coded into each theme.

| Code | Example quote |
| --- | --- |
| 'Explicit positive change' [starting with a 'yes'] | *"Yes, I especially notices [sic] how young the majority of them were, which surprised me in a good way."* |
| 'Explicit positive change' [without a yes] | *"Slightly. Females are a part of science/biology more than we realise."* |
| 'Explicit no change' [starting with a 'no'] | *"No - I don't understand the importance or relevance of having academic headshots on lecture slides - we have the name and study if we want to look into them further but we have no need to know what they look like."* |
| 'Explicit no change' [without a 'no'] | *"It has not changed"* |
| Ambivalence | *"Not going to lie I was too busy laughing at the first guy called Pu Tang"* |
| Diversity Assumed | *"No, I already had the mindset that it would be a wide range of people carrying out the research."* |

The final thematic analysis process was for responses to Question 8 - “Do you feel this is good practice for lectures (please elaborate)?”. Similarly to the previous question, most answers began with a ‘Yes’ or ‘No’ response, which provided the first level of coding. ‘Yes’ responses were automatically coded into the theme ‘Explicit good practice’, while ‘No’ responses into ‘Explicit not good practice’. In a few cases, this was the whole of the answer. Not all questions started with a ‘Yes’/’No’ statement, in these instances the response was reviewed for value-statements that were inherently positive or negative. Responses that were deemed to be unambiguously positive were coded to Explicitly good practice, while those that were negative were coded to ‘Explicit not good practice’. These were generally clear but conversations with the second coder were used to ensure consistency of response. The final code allocated was Ambivalent, which was linked to responses that highlighted both positive and negative elements, anyone who was unsure of the impact, or a response that was unrelated to the question. Once the highest level themes were coded each was reviewed separately for similarities in the rationale for participant answers. These sub-themes tied together the different types of reason the participants had given for answering the way that they had. Responses could be coded into multiple sub-themes, particularly where the open text response was long and comprehensive. The sub-themes that arose are considered in detail in the results section.

**Supplementary Table C.** Examples of responses to a question asking whether the participant felt that intervention was good practice, focusing particularly on how responses were coded into each theme.

| Code | Example quote |
| --- | --- |
| 'Explicit good practice' [starting with a 'yes'] | *"Yes, it's nice to see people like me!"* |
| 'Explicit good practice' [without a yes] | *"Good idea helps to picture the person and their experiences"* |
| 'Explicit not good practice' [starting with a 'no'] | *"No, prefer to have the links to the paper so I can read them if I want."* |
| 'Explicit good practice' [without a 'no'] | *"I would rather focus on the information rather than the author"* |
| Ambivalence | *"Interesting but unsure of relevance"* |

